# Robust Tobacco Smoking Self-Report in two Cohorts of Vulnerable Pregnant Women and Adults

**DOI:** 10.1101/2020.09.11.293902

**Authors:** Sara Saberi, Marie-Soleil R. Smith, Abhinav Ajaykumar, Mayanne M. T. Zhu, Izabelle Gadawski, Beheroze Sattha, Evelyn J. Maan, Julie Van Schalkwyk, Chelsea Elwood, Isabelle Boucoiran, Deborah M. Money, Hélène C.F. Côté, CIHR Team in Cellular Aging and HIV Comorbidities in Women and Children (CARMA)

## Abstract

**Background:** Stigma associated with tobacco smoking, especially during pregnancy, may lead to underreporting and possible bias in studies relying on self-reported smoking data. Cotinine, a nicotine metabolite with a ∼20h half-life in blood, is often used as a biomarker of smoking. The objective of this study was to examine the concordance between self-reported smoking and plasma cotinine concentration among participants enrolled in two related cohorts of vulnerable individuals: human immunodeficiency virus (HIV)-positive and HIV-negative pregnant women enrolled in the CARMA-PREG cohort and HIV-positive and HIV-negative non-pregnant women and men enrolled in the CARMA-CORE cohort.

**Methods:** For HIV-positive (n=76) and negative (n=24) pregnant women, plasma cotinine was measured by ELISA in specimens collected during the third trimester, between 28 and 38 weeks of gestation. Plasma cotinine was also measured in HIV-positive (n=43) and negative (n=57) women and men enrolled in the CARMA-CORE cohort.

**Results:** Self-reported smokers were more likely to have low income (p<0.001) in both cohorts, and to deliver preterm (p=0.007) in CARMA-PREG. In the CARMA-PREG cohort, concordance between plasma cotinine was 95% for self-reported smoking, and 89% for self-reported non-smoking. In the CARMA-CORE cohort we observed similarly high concordances of 96% and 92% for self-reported smoking and non-smoking, respectively. In this sample, the odds of discordance between self-reported smoking status and cotinine levels were not significantly different between self-reported smokers and non-smokers, nor between pregnant women and others. Taken together, the overall concordance between plasma cotinine and self-reported data was 94% with a Cohen’s kappa coefficient of 0.860 among all participants.

**Conclusions:** Given the high proportion of vulnerable people in the CARMA-PREG and CARMA-CORE cohorts, our results may not be fully generalizable to the general population. However, they demonstrate that participant surveying in a non-judgemental context can lead to accurate and robust self-report data.

**Implications:** Reliable self-reported smoking data is necessary to account for smoking status in subsequent studies. Our results suggest that future studies should ensure that study participants feel sale to speak candidly to non-judgemental research staff to obtain reliable self-report data.

## Introduction

The adverse health effects of tobacco smoking are well-documented ^1^ and smoking is a well-established risk factor for adverse pregnancy outcomes, including the likelihood of preterm delivery, intrauterine growth retardation, and placenta previa ^2-7^. Prenatal tobacco exposure is also associated with low birth weight, higher risk of sudden infant death syndrome, and adverse health outcomes later in life ^3^. Most cohort studies collect information on smoking using self-report data; however, depending on the social context, the stigma associated with smoking, especially during pregnancy, may lead to underreporting of this behavior and possible bias in studies relying on self-reported smoking data ^8-12^. Indeed, previous studies reported higher non-disclosure rates of smoking among pregnant women than the general population ^13-15^. For example, in a smoking cessation trial in the USA, pregnant women underreported their smoking by 14% ^13^, and an Australian study also reported that 7–15% of pregnant women did not disclose their smoking habit ^11^.

In 2016, over 1.1 billion (21.9%) people smoked tobacco, varying widely from country to country from 4% to 47% ^16^. The rate of smoking among pregnant women also varies around the world, from 10% in Japan ^17^ to 17% in Australia ^18^ and 30-35% in Spain ^19^. Approximately one in five Canadians, or 5.8 million people, reported smoking in 2011 ^20^. Among pregnant women, the prevalence of smoking was estimated to be 17% in 2000-01 ^21^, although prevalence data are limited and rates vary greatly between regions ^21-23^. Data is also scarce on the accuracy of self-reported smoking behaviors during pregnancy.

Cotinine, a nicotine metabolite with a ∼16h half-life in plasma, urine, or saliva, is often used as a biomarker of smoking and can be used to determine concordance between levels of the marker to self-reported smoking status ^24^. Cotinine has also been identified in cord blood and infant urine as an indicator of fetal exposure to tobacco ^25^. The objective of this study was to examine the concordance between self-reported smoking and plasma cotinine concentration among HIV-positive (HIV+) and HIV-negative (HIV-) pregnant women enrolled in the Canadian CARMA (Children and women, Antiretrovirals and Markers of Aging)-PREG Cohort, HIV+ and HIV-women and men enrolled in the Canadian CARMA-CORE Cohort, and HIV.

## Methods

CARMA-PREG and CARMA-CORE are prospective cohort studies of HIV+ and HIV-pregnant women enrolled between 2004 and 2020, and HIV+ and HIV-non-pregnant women and men enrolled in 2008 to 2017, respectively. Inclusion criteria for enrolment in the CARMA-PREG cohort were being pregnant with a known HIV status, in which those who were HIV+ women had to be receiving or be willing to receive antiretroviral therapy during pregnancy. Enrolment in CARMA-CORE took place during routine clinical visits at four sites across Canada and HIV-controls were recruited to have similar sociodemographic characteristics as the HIV+ group.

For CARMA-PREG, HIV+ women were recruited from the Oak Tree Clinic at British Columbia Women’s Hospital and the Sainte-Justine Hospital in Quebec. HIV-participants were recruited through a variety of means, including word of mouth and advertisements posted in strategic areas in Vancouver, British Columbia to promote enrollment of HIV-control women with similar sociodemographic characteristics to HIV+ participants, including smoking habits. The enrolment of women in CARMA-PREG took place in the first trimester and both HIV+ and HIV-women provided biological specimens at three visits during pregnancy and at delivery. Exclusion criteria for the cohorts included the inability to provide informed consent (language barriers) or to participate in research (health or social crisis). Additional exclusion criteria for the selection of participants included the exclusion of those missing 3^rd^ visit plasma specimens for CARMA-PREG and missing 1^st^ visit plasma specimens for CARMA-CORE, those with missing smoking information due to incomplete data collection or refusal to answer, and those enrolled more than once in a CARMA cohort (repeat pregnancies in which any additional pregnancies were excluded, or enrolment in both CARMA-PREG and CARMA-CORE in which the CARMA-PREG data was prioritized) (Figure S1). Fifty-seven non-smoking participants in CARMA-PREG were selected to balance year of visit with the remaining n=43 smokers, for a total of 100 participants. Fifty smokers and fifty non-smokers in CARMA-CORE were sex balanced and then randomly selected for a total of 100 participants.

For both cohorts at each visit demographic, clinical, and substance use information, including tobacco exposure, were collected by self-report (any tobacco use since last visit). At the time of data collection, tobacco exposure occurred exclusively through cigarette smoking and not vaping. One CARMA-PREG participant included in this study reported chewing tobacco at a rate equivalent to a pack of cigarettes a day and was treated in this analysis as such. No data were ever collected about second-hand smoking or nicotine patch or gum use. Participants were not informed that smoking status would be confirmed with cotinine concentrations, thus data were anonymized.

Plasma cotinine levels were measured in specimens collected at the 3^rd^ visit between 28 and 38 weeks of gestation from a total of 76 HIV+ and 24 HIV-CARMA-PREG participants. Additional plasma specimens at delivery and cord blood from select participants, 7 with high cotinine values and 7 with discordant self-report and cotinine values, were also assessed. In the CARMA-CORE cohort, plasma cotinine levels were measured in specimens collected during their first visit from a total of 43 HIV+ and 57 HIV - participants. Self-reported substance use information was collected on the same day as blood collection. Cotinine was measured by solid phase competitive enzyme-linked immunosorbent assay (ELISA) (Calbiotech, CA, USA).

Comparisons between self-reported smokers and non-smoker groups were done using Fisher’s exact test or Chi-square for categorical variables, and Student t-test for continuous variables. Defining smoking by a plasma cotinine ≥5 ng/ml, the proportions of cotinine-negative among self-reported non-users and of cotinine-positive among self-reported users were used to express concordance. In addition, Cohen’s kappa coefficient (κ) was used to determine the agreement between smoking as per self-report and cotinine among all participants and within the HIV+ and HIV-groups. All statistical analyses were performed using GraphPad Prism 8.4.3.

## Results

In this study, 43 (43%) of CARMA-PREG participants self-reported smoking during pregnancy between 28 and 38 weeks of gestation, representing the high rate of smoking in CARMA-PREG cohort (Table 1). There was no significant difference in maternal age between the self-reported smokers and non-smokers, but HIV status was more prevalent among self-reported smokers (p=0.04). Eighty-six (86)% of self-reported smokers and 68% of self-reported non-smokers were living with HIV. There were fewer Indigenous/First Nations women in the non-smokers group and fewer African Caribbean Black women in the smokers group (p<0.001). Compared to non-smokers, self-reported smokers were more likely to have a preterm delivery (<37 weeks of gestation) (p=0.007) to have a low income (<$15000/year, p<0.001), and to use illicit drugs (p<0.001) (Table 1).

**Table 1.**
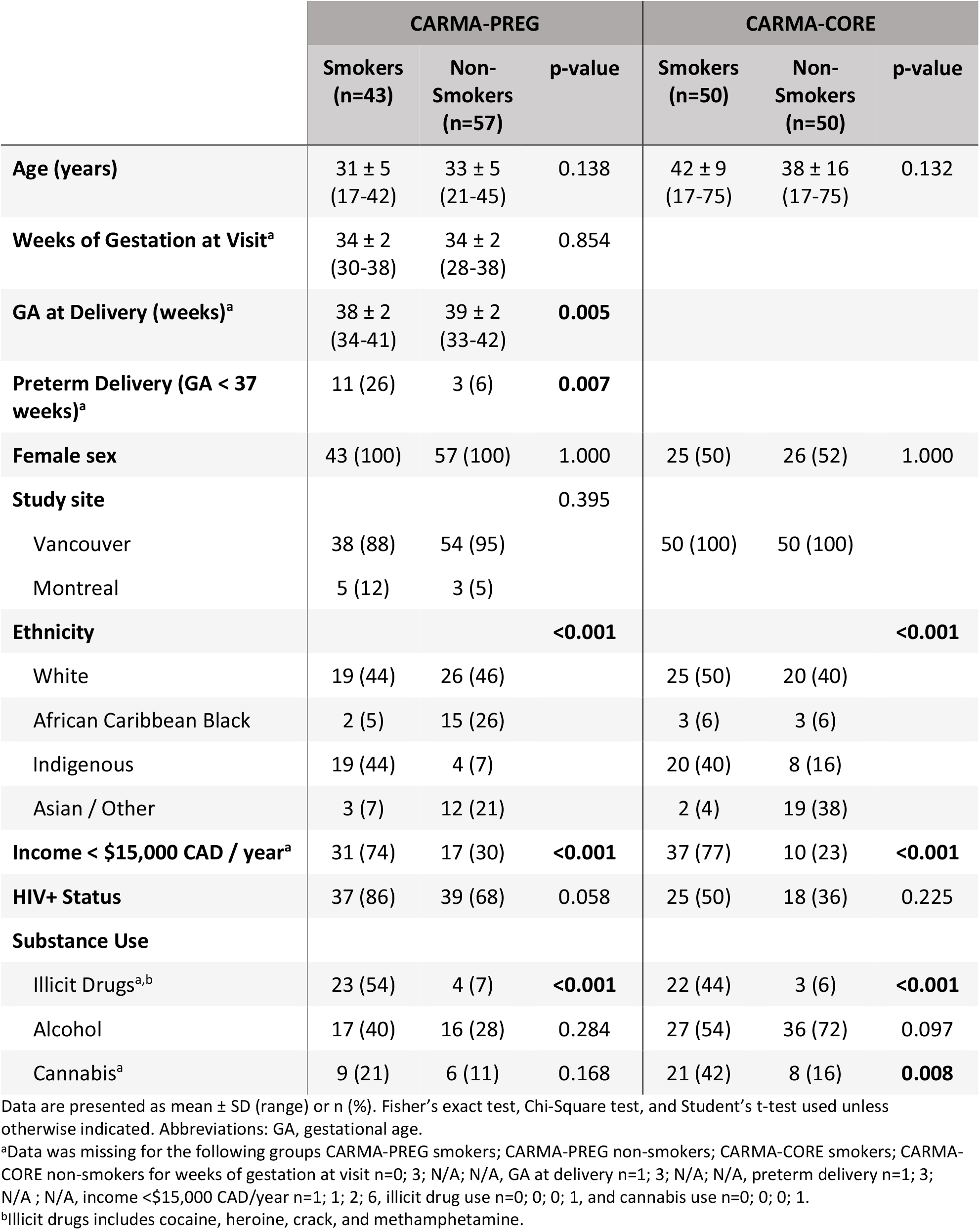
Demographic, clinical, and substance use characteristics of the study participants from two separate cohorts self-reporting tobacco smoking at their study visit, or not.

**Table 2.**
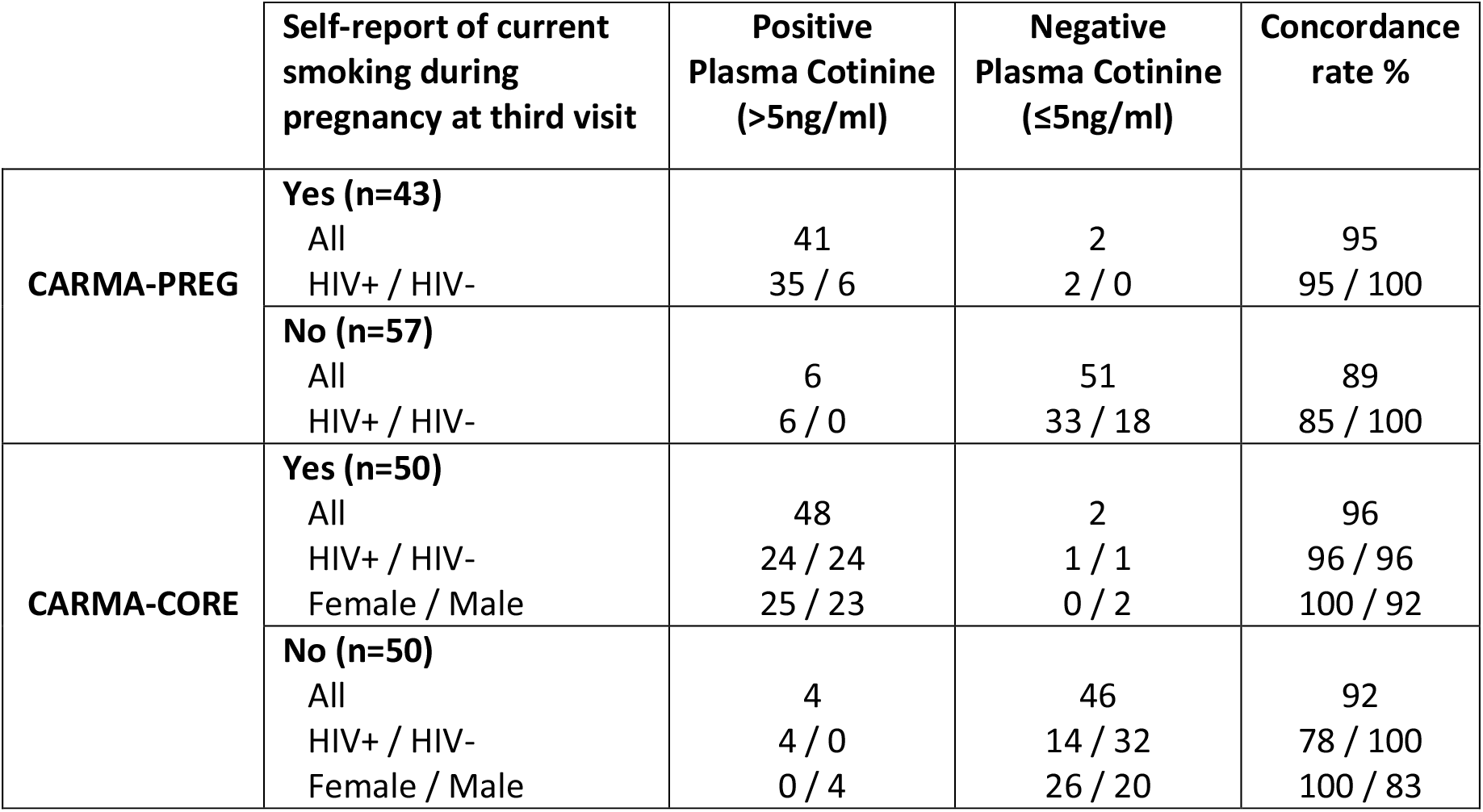
Concordance between self-reported smoking at study visit and plasma cotinine concentration measured by ELISA on a plasma specimen collected day of visit.

|  | Self-report of current smoking during pregnancy at third visit | Positive Plasma Cotinine (>5ng/ml) | Negative Plasma Cotinine (≤5ng/ml) | Concordance rate % |
| --- | --- | --- | --- | --- |
| **CARMA-PREG** | **Yes (n=43)**<br>All<br>HIV+ / HIV- | 41<br>35 / 6 | 2<br>2 / 0 | 95<br>95 / 100 |
|  | **No (n=57)**<br>All<br>HIV+ / HIV- | 6<br>6 / 0 | 51<br>33 / 18 | 89<br>85 / 100 |
| **CARMA-CORE** | **Yes (n=50)**<br>All<br>HIV+ / HIV-<br>Female / Male | 48<br>24 / 24<br>25 / 23 | 2<br>1 / 1<br>0 / 2 | 96<br>96 / 96<br>100 / 92 |
|  | **No (n=50)**<br>All<br>HIV+ / HIV-<br>Female / Male | 4<br>4 / 0<br>0 / 4 | 46<br>14 / 32<br>26 / 20 | 92<br>78 / 100<br>100 / 83 |

Among women who reported having smoked (n=43) since their last visit during pregnancy in the CARMA-PREG cohort, 3 (7%) reported smoking heavily (a pack a day or more), 28 (65%) reported smoking a moderate amount (2 to 19 cigarettes a day), and 3 (7%) reported light smoking (less than 2 cigarettes a day). The frequency and/or quantity of tobacco use was unavailable for the remaining 9 (21%). Defining smoking by a plasma cotinine ≥5 ng/mL, we observed 95% concordance between self-reported smoking and plasma cotinine. For the women who self-reported as non-smokers, concordance with plasma cotinine <5 ng/mL was 89%. Two pregnant women who self-reported smoking showed plasma cotinine levels <5 ng/ml. According to their smoking frequency data, both women reported smoking on average fewer than two cigarettes per week during their pregnancy. The κ for women in CARMA-PREG was 0.839, indicating almost perfect agreement. Among pregnant women living with HIV (n=76), the concordance between plasma cotinine and self-reported smoking (n=37) and non-smoking (n=39) was 95% and 85%, respectively. Among pregnant women not living with HIV (n=24), 6 self-reported smoking and the remaining 18 reported no smoking, and the observed concordance was 100%. The κ for HIV+ and HIV-pregnant women, were 0.790 and 1.00, respectively.

To further examine the quality of self-reported smoking data in our cohorts we examined the concordance of self-reported smoking data and plasma cotinine in the CARMA-CORE cohort. Among the women and men who reported tobacco use at their first visit in the CARMA-CORE cohort, 11 (22%) reported smoking heavily (a pack a day or more), 29 (58%) reported smoking a moderate amount (2 to 19 cigarettes a day), and 7 (14%) reported light smoking (less than 2 cigarettes a day). The frequency and/or quantity of tobacco use was unavailable for the remaining 3 (6%). We observed 96% concordance between self-reported smoking and plasma cotinine. For the women and men who self-reported as non-smokers, concordance with plasma cotinine <5 ng/mL was 92%. The κ for participants in CARMA-CORE was 0.880. Among all CARMA-CORE participants living with HIV in this study (n=43), the concordance between plasma cotinine and self-reported smoking (n=25) and non-smoking (n=18) was 96% and 78%, respectively. Among the women and men not living with HIV (n=57), 25 self-reported smoking and the remaining 32 reported no smoking, and the observed concordance was 96% and 100%, respectively. The κ for HIV+ and HIV-CARMA-CORE participants were 0.755 and 0.964, respectively. Among all female CARMA-CORE participants (n=51), the concordance between plasma cotinine and self-reported smoking (n=25) and non-smoking (n=26) was 100%. Male CARMA-CORE participants (n=49), had a concordance between plasma cotinine and self-reported smoking (n=25) and non-smoking (n=24) of 92% and 83%, respectively. The κ for female and male CARMA-CORE participants were 1.00 and 0.755, respectively.

Taken together, if we group the two datasets, among 100 pregnant women, 51 non-pregnant women, and 49 men included in our analyses, 93 (47%) self-reported smoking and we observed an overall concordance of 94% between plasma cotinine and both self-reported smoking and non-smoking data with a κ of 0.860. Among all individuals living with HIV in both cohorts (n=119), the concordance between plasma cotinine and self-reported smoking (n=62) and non-smoking (n=57) was 95% and 81%, respectively. Among all individuals who did not have an HIV diagnosis (n=81), 31 self-reported smoking and the remaining 50 reported no smoking, with an observed concordance of 98% and 100% respectively. The κ for HIV+ and HIV-individuals were 0.780 and 0.974 respectively. Among all women in both cohorts (n=151), the concordance between plasma cotinine and self-reported smoking (n=68) and non-smoking (n=83) was 98% and 95%, respectively, with a κ of 0.894.

We also investigated the relationship between cotinine concentrations from seven self-reported smokers with high cotinine and seven discordant self-report participants in CARMA-PREG in plasma at third visit and plasma and cord plasma at delivery. Participants who reported smoking and had high cotinine levels at 3^rd^ visit also had high levels of cotinine in maternal and cord plasma at delivery (Figure S2). Two of the seven participants with discordant self-report and cotinine values reported light smoking but consistently had cotinine values <5 (Figure S3). Of the five remaining discordant participants who self-reported non-smoking but had cotinine values >5 at third visit, three had cotinine values <5 at delivery in maternal and cord plasma and two remained >5 throughout (Figure S3).

For quality control purposes, we repeated the ELISA on 74 of the CARMA-PREG specimens. Qualitatively, we obtained identical results. Quantitatively, the plasma cotinine concentrations measured at two different times were strongly correlated (r=0.601, p<0.001) indicating the assay is robust enough to assay specimens only once (Figure S4).

## Discussion

The prevalence of maternal smoking in the CARMA-PREG cohort (43%) is higher than the estimated rate of smoking during pregnancy in Canada ^6,21,26^, but is similar to reported rates among HIV+ persons in Canada ^27^. Although the rates of smoking during pregnancy in our CARMA-PREG cohort are higher than their corresponding regions in Canada, they are comparable to that of pregnant women in the Canadian Northern Territories (39.4 in 2005 and 59.3% in 2010) ^6,26^, which is likely related to the ethnic makeup and socioeconomic status of our cohort. Although an even selection of smokers and non-smokers were selected from the CARMA-CORE cohort for cotinine analysis, the rate of smoking in the entire cohort in which smoking info was available (n=623) was 40%, twice as high as the Canadian average, but again similar to reported rates among HIV+ persons in Canada ^20,27^. Approximately 44% of participants who self-reported smoking in CARMA-PREG cohort and 40% in CARMA-CORE were Indigenous. Additionally, our results are consistent with previous Canadian studies showing that low income is associated with higher smoking rate during pregnancy ^6,21,26^. These observations suggest that socio-cultural and socio– economic determinants rather than place of residence may be associated with smoking rates.

Many studies reporting smoking data rely on self-report. The CARMA cohorts use this same method, but we additionally undertook the current study to examine the concordance between self-report and a marker of smoking, namely plasma cotinine levels. In participants from two separate cohorts we observed excellent concordance between the two, both among HIV+ and HIV-participants and in the pregnancy and non-pregnancy cohorts. Only 9% of self-reported non-smoker participants were likely to be smoker considering their cotinine test result. In addition, 4% of self-reported smokers showed plasma cotinine levels ≤5ng/ml, but most were participants who reported light smoking intensity. Given the half-life of cotinine, the timing of maternal sampling could influence its detection. Hence, lower concordance between cotinine and self-reported smoking may be expected using this assay when the smoking frequency is very low.

Our results agree with a Swedish study reporting that 6% of self-reported non-smoker pregnant women were more likely to be smokers and that 3% had cotinine concentrations suggestive of passive smoking ^28^. However, many other studies report lower concordance. For example, in a study taking place in West Scotland, 25% of pregnant women who self-reported to be non-smokers had cotinine measurements above the threshold ^11^. In rural and small metropolitan areas in upstate New York, 35% of women who self-reported not smoking during pregnancy, had urinary cotinine levels suggesting inaccurate self-reported status ^9^. Such cases of underreporting may introduce bias to the associated measures and may have implications not only in the research setting but for clinicians and caregivers who assess past and present tobacco use ^10^. The integrity of self-reported data varies according to population and the social context in which the data are collected. Multiple factors could influence the interviewee’s responses to questions about smoking status, including participant characteristics, study method and setting, and the pressure mediated by social desirability.

A strength of this study is the high smoking rate within the cohorts, among both HIV+ and HIV-participants and the multi-ethnic makeup. Although the cohort may not be fully generalizable to pregnant and non-pregnant populations, our results showed high concordance among both HIV+ (90%) and HIV- (100%) pregnant women, HIV+ (96%) and HIV- (96%) non-pregnant women and men. A limitation to this study is that the intensity of smoking was not recorded for all participants, which prevented more detailed analyses. Another limitation is the absence of self-reporting on nicotine patch or gum use, which may help explain some of our discordant results ^29^.

Overall, our data suggest that it is possible to obtain robust data on smoking from pregnant and non-pregnant participants through self-report. Among CARMA cohort participants, our results indicate that self-reported smoking data are highly reliable as a surrogate for tobacco exposure. This suggests that study participants likely felt safe to speak candidly and accurately self-reported their smoking habits to our non-judgemental research staff.

## Acknowledgements

The authors report no conflict of interest for this study. This research was supported in part by two Canadian Foundation for AIDS Research grants (016012 to DMM & HCFC, and 0120 004 to HCFC & DMM) as well as two Canadian Institutes of Health Research (CIHR) team grants (HET-85515 to HCFC & DMM, and TCO-125269 to HCFC & DMM). SS (2012-13) and AA (2016-17) were partially supported by University of British Columbia (UBC) Centre for Blood Research Collaborative training awards. AA was also supported partially by two UBC Faculty of Medicine scholarships and MSRS was also partially supported by the British Columbia Graduate Student Award. We thank all study participants and the dedicated Oak Tree Clinic research staff and students including Rebecca Graham, Ashley Docherty, Jackson Chu, Daljeet Mahal, Laura Oliveira, Farnaz Changizi, Tessa Chaworth-Musters, Maninder Mahal, Annie Qiu, Clare Hall-Patch, Ariel Nesbitt, and Shanlea Gordon.

## Author’s contributions

Marie-Soleil Smith: Assisted in the study design, assayed (ELISA) specimens, performed the analysis, and contributed to writing the manuscript.

Sara Saberi: Assisted in the study design, assayed (ELISA) specimens, assisted with basic statistical analysis, and contributed to writing the manuscript.

Abhinav Ajaykumar: Participated in the study design and assisted in drafting the manuscript. Mayanne M. T. Zhu: Assisted with sample preparation and basic statistical analysis; reviewed the final manuscript.

Izabelle Gadawski: Assisted with ELISA assay and reviewed the manuscript.

Beheroze Sattha: Assisted with sample preparation, data retrieval, and reviewed the manuscript. Evelyn Maan: Assisted with HIV+ and HIV-participant enrollment and data collection. Contributed to database building, critically reviewed the manuscript.

Julie Van Schalkwyk: Contributed to enrollment of HIV-participants and critically reviewed and edited the manuscript

Deborah M. Money: CARMA-PREG principal investigator (PI), contributed to its study design, enrollment of HIV+ participants, and critically reviewed and edited the manuscript.

Hélène C.F. Côté: CARMA cohort study PI, designed the study, critically reviewed, and edited the manuscript

## Ethical considerations

Ethical approval for this secondary use of data study was obtained from the Research Ethics Boards of the University of British Columbia and from the Hospital Research Review Committee of the Children’s and Women’s Health Centre of British Columbia (H03-70356, H04-70540, H07-03136, and H08-02018). All participants provided written informed consent to have their cohort specimens biobanked.

---

**Figure 1.**
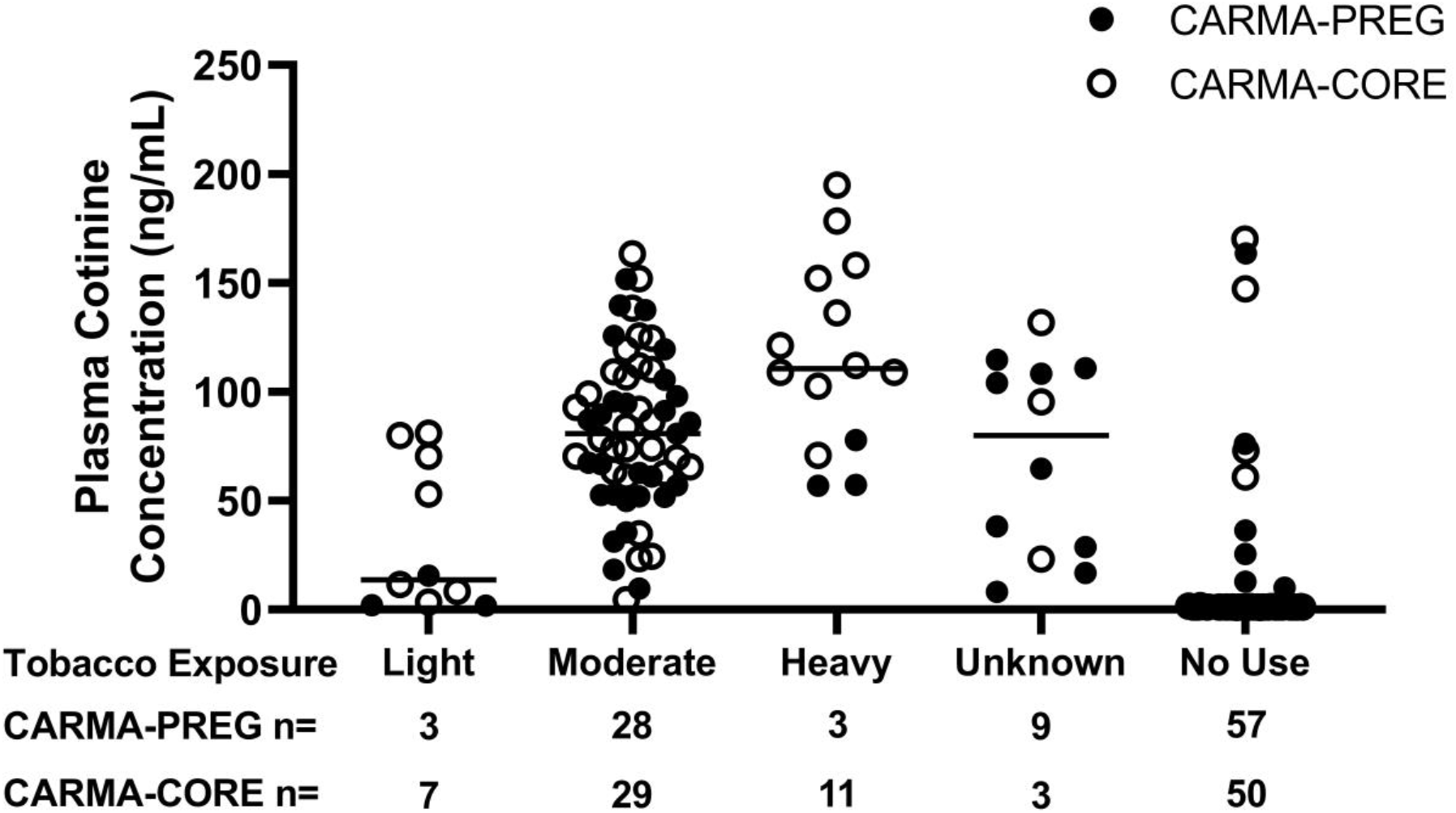
Plasma cotinine concentration according to self-reported intensity of smoking since last visit. Median cotinine are represented by the horizontal bar.

**Figure S1.**
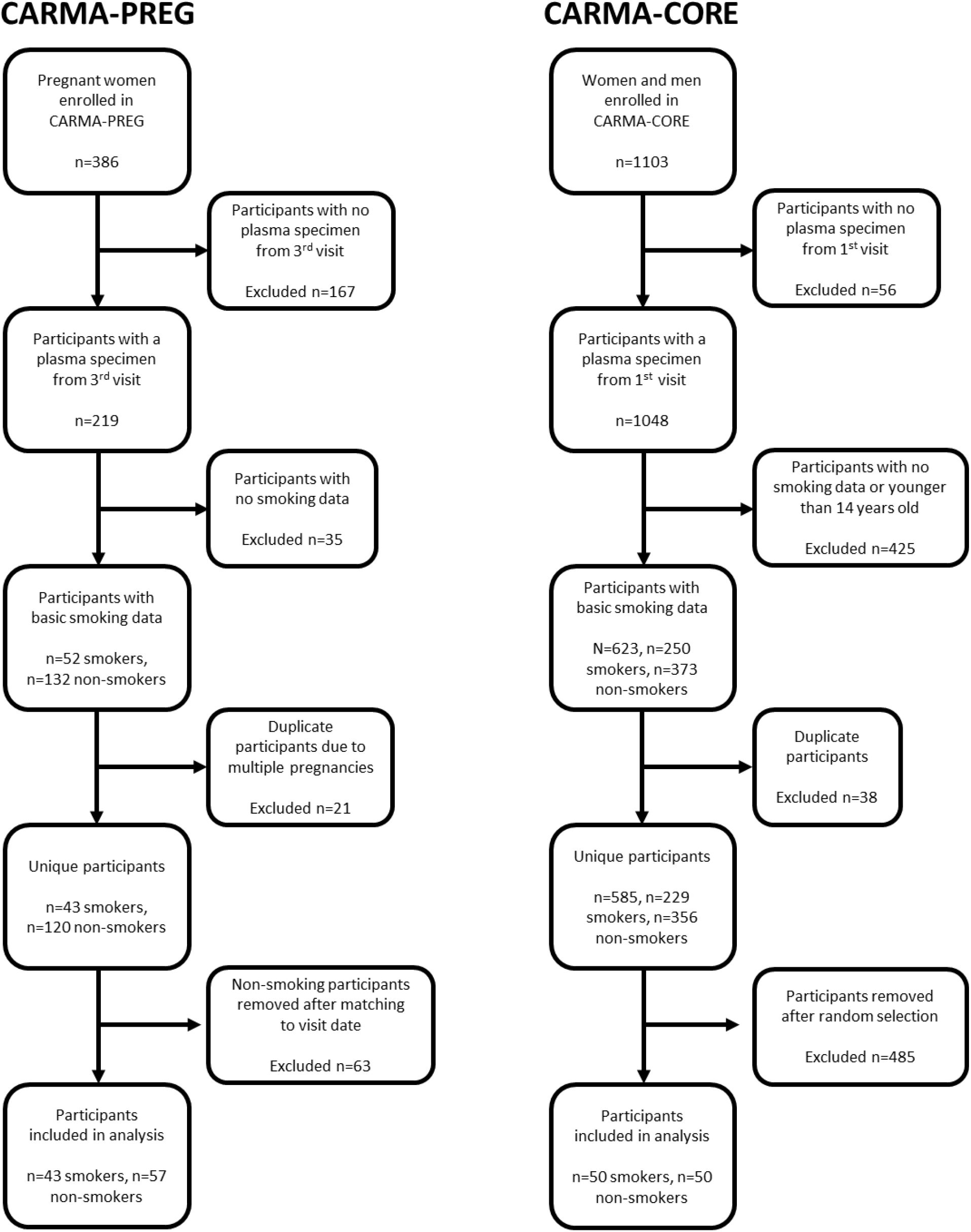
Selection criteria for both cohorts.

**Figure S2.**
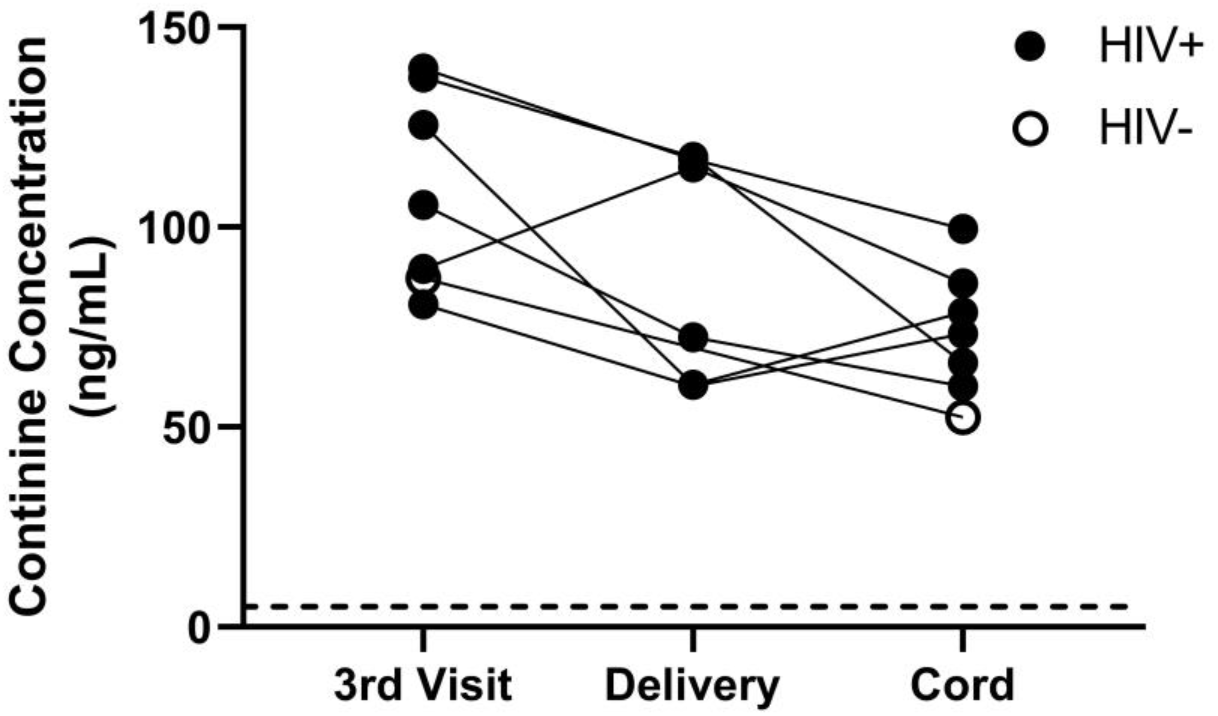
Cotinine concentrations from 7 self-reported smokers with high cotinine in plasma at 3^rd^ visit and plasma (n=6) and cord plasma (n=7) at delivery. Dotted line at y=5 indicates cotinine positivity.

**Figure S3.**
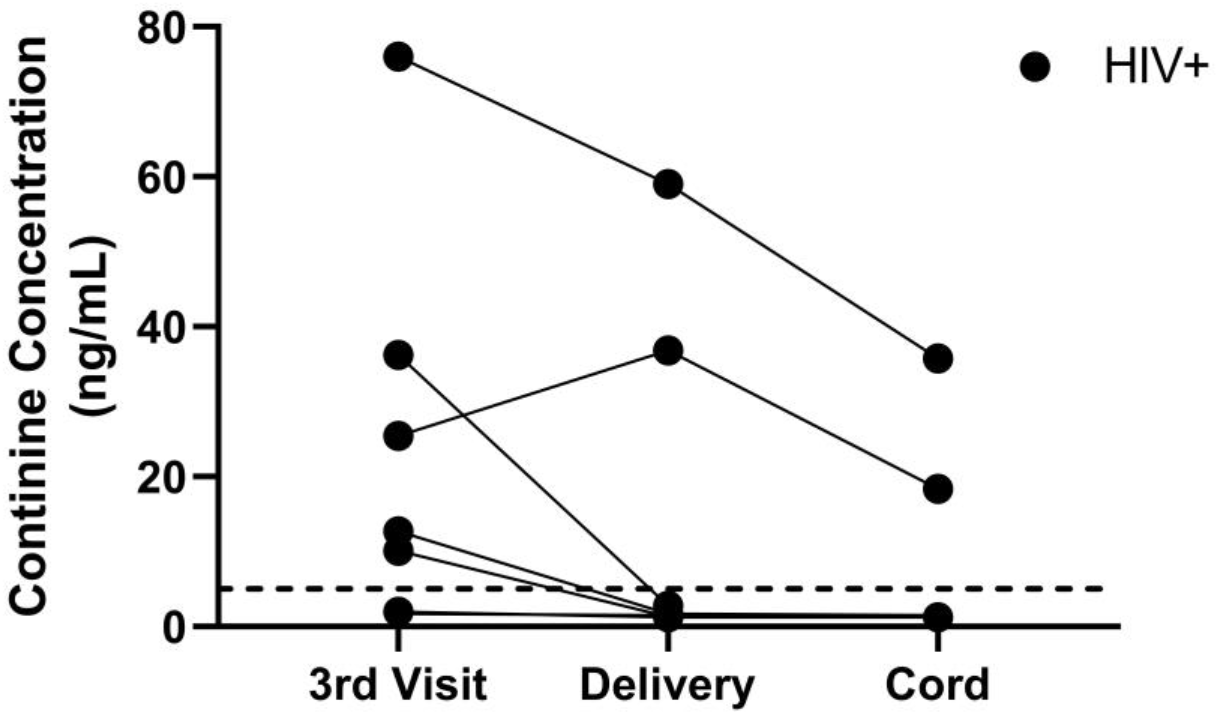
Cotinine concentrations from 7 discordant self-report participants in plasma at 3^rd^ visit and plasma (n=6) and cord plasma (n=6) at delivery. Dotted line at y=5 indicates cotinine positivity.

**Figure S4.**
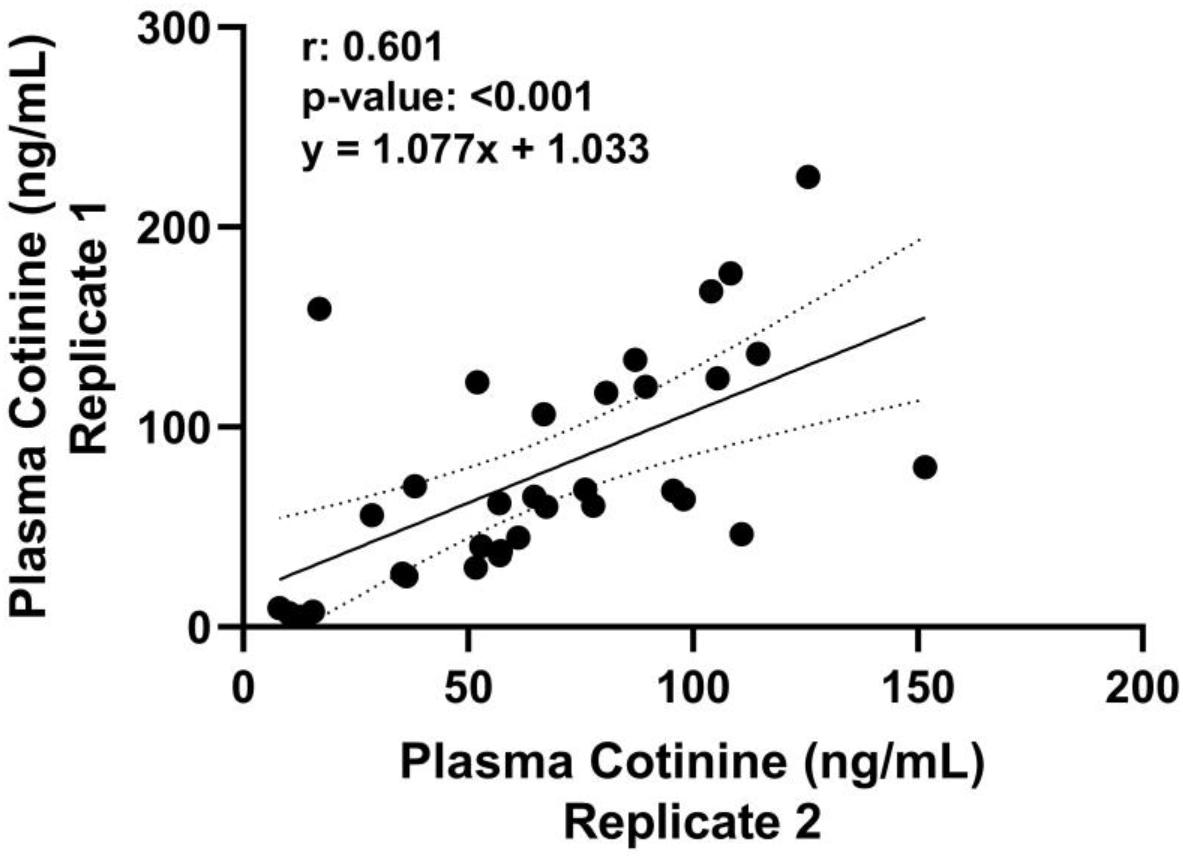
Correlation between n=74 CARMA-PREG specimens assayed once in 2016 (Replicate 1) and in 2020 (Replicate 2). Lines indicate best fit and 95% confidence intervals.

